# Development and Characterization of a 2D Porcine Colonic Organoid Model for Studying Intestinal Physiology and Barrier Function

**DOI:** 10.1101/2024.10.18.619022

**Authors:** Masina Plenge, Nadine Schnepel, Mathias Müsken, Judith Rohde, Ralph Goethe, Gerhard Breves, Gemma Mazzuoli-Weber, Pascal Benz

## Abstract

The porcine colon epithelium plays a crucial role in nutrient absorption, ion transport, and barrier function. However ethical concerns necessitate the development of alternatives to animal models for its study. The objective of this study was to develop and characterise a two-dimensional (2D) *in vitro* model of porcine colonic organoids that closely mimics native colon tissue, thereby supporting in vitro research in gastrointestinal physiology, pathology, and pharmacology. Porcine colonic crypts were isolated and cultured in three-dimensional (3D) organoid systems, which were subsequently disaggregated to form 2D monolayers on transwell inserts. The integrity of the monolayers was evaluated through the measurement of transepithelial electrical resistance (TEER) and electron microscopy. The functional prerequisites of the model were evaluated through the measurement of the mRNA expression of key ion channels and transporters, using quantitative RT-PCR. Ussing chamber experiments were performed to verify physiological activity. The 2D monolayer displayed robust TEER values and retained structural characteristics, including microvilli and mucus-secreting goblet cells, comparable to those observed in native colon tissue. Gene expression analysis revealed no significant differences between the 2D organoid model and native tissue with regard to critical transporters. Ussing chamber experiments demonstrated physiological responses that were consistent with those observed in native colonic tissue. In conclusion, 2D porcine colonic organoid model can be recommended as an accurate representation of the physiological and functional attributes of the native colon epithelium. This model offers a valuable tool for investigating intestinal barrier properties, ion transport, and the pathophysiology of gastrointestinal diseases, while adhering to the 3R principles.

## Introduction

The gastrointestinal tract (GIT) is vital for nutrient digestion and absorption, waste excretion, and metabolic homeostasis (1). The GIT epithelium acts as a selective barrier, regulating the passage of nutrients, ions, and water. Tight junction proteins, such as occludin (OCLN), claudin (CLDN) and zonula occludens (ZO-1), are crucial in maintaining the integrity of the epithelial barrier by providing both a “fence” and a “gate” or barrier function (2, 3). The fence function serves to separate the apical from the basolateral sides of the epithelial cells, ensuring the proper distribution of membrane proteins and lipids. Moreover, the mucus layer, which is composed of mucins (MUC) and is secreted by goblet cells, serves as the primary defence against luminal pathogens and mechanical stress (4). Various transporters, channels, and carriers facilitate the transport across the epithelium, such as the epithelial sodium channel (ENaC), sodium-hydrogen antiporter (NHE), calcium-dependent chloride channel (CaCC), and cystic fibrosis transmembrane conductance regulator (CFTR). Each of these regulates the transport of specific ions and molecules. The function of the GIT has historically been studied by animal experiments, with pigs being commonly used as models due to their relevance to both, pig-specific diseases and human intestinal conditions (5).

However, with the increasing relevance of ethical concerns and the principles of replacement, reduction, and refinement (3R) (6), alternative methodologies such as cell culture-based systems are gaining attraction. Cell lines have been instrumental in unravelling certain aspects of GIT physiology. Nevertheless, the usefulness of these models has often been limited by their focus on singular cell types. This limitation highlights the need for more comprehensive models that encompass the diverse cell populations found within the gastrointestinal epithelium. The colon hosts a variety of cell types, including enterocytes, goblet cells, enteroendocrine cells, tuft cells, and stem cells, each contributing uniquely to its function (7, 8). To overcome this limitation of a single cell type, the organoid model was introduced in 2009 (9), revolutionising our ability to model the complex architecture and functionality of the intestinal epithelium. Organoids are derived from stem cells, which impart two crucial properties to them. Firstly, these cells can differentiate into all cell types of the epithelium from which they were obtained. Secondly, organoids possess the ability to self-renewal (10). Organoids cultivated in a 2D-monolayer system are highly valued as effective tools for studying barrier properties for drug discovery and development (11). According to Hoffmann et al. (12), porcine jejunum organoid cultures are a useful model for investigating physiological transport properties of the epithelium. Nevertheless, a comprehensive model that takes into account the physiological attributes of the porcine colon is still lacking. Colon organoids offer unique advantages for investigating colon-specific disease mechanisms and treatments. They effectively model the thicker mucus layer that is essential for maintaining epithelial barrier integrity, and facilitate research into critical functions such as water and electrolyte absorption. Thus, this study aimed to delineate the physiological functionality of a 2D culture of porcine colonic organoids, emphasizing gene expression relative to native tissue and elucidating the transport properties of the organoid-derived epithelium.

## Material & methods

### Culture of 3D-organoids

To generate 3D organoids, intestinal crypts were isolated from the porcine colon. The cultivation of 3D organoids has already been described in detail by Hoffmann et al. (12). Briefly, the organoids were cultured in Matrigel (Corning®, Kaiserslautern, Germany; Matrigel® Basement Membrane Matrix) droplets á 50 µl in a 24-well plate. Every 2 – 3 days, the medium was changed and at least once a week the droplet was resolved and the 3D organoids were passaged. Thereby the medium (S1 Table) was aspirated, ice-cold PBS added and the Matrigel droplet dissolved by pipetting. The suspension was collected in a tube and was centrifuged at 500 x *g* for 10 minutes at 4 °C. The pellet was resuspended in medium and diluted 1:5 with Matrigel. The plate was incubated at 37 °C for at least 30 min and overlaid with 500 µl of medium per well.

### Culture of 2D-organoids

2D organoids were generated by enzymatic and mechanical disintegration of 3D organoids as described in the following. The medium was aspirated and the Matrigel droplets were resolved with 1 ml ice cold PBS, at this stage 3 – 4 wells were pooled. The solution was transferred to a reaction tube with 10 ml ice cold PBS and then centrifuged at 250 x *g* for 10 min at 4 °C. The supernatant was removed, the pellet was resuspended in 1 ml 0.05 % Trypsin/ EDTA (Gibco™, Thermo Fisher) and incubated for 5 min at 37 °C. Afterwards, the solution was resuspended 20 times with a 1,000 µl tip and 15 times with a 200 µl tip, which was mounted on a 1,000 µl tip, all steps being performed on ice. 10 ml Advanced DMEM/F-12 (Thermo Fisher scientific, Waltham, USA) supplemented with 10% (v/v) fetal bovine serum (FBS; (Sigma-Aldrich, Darmstadt, Germany; catalogue no.: F7524) were added and centrifuged at 1,000 x *g* for 10 min at 4 °C. After removing the supernatant, the pellet was resuspended in 1 ml monolayer medium (S2 Table). Cells were counted and 1.5 ‧ 10^5^ cells were seeded on cell culture inserts (Corning®; Snapwells® or GREINER BIO ONE; ThinCert® Cell Culture Insert 12 Well; diameter: 12 mm; pore size: 0.4 μm) which were pre-coated with 1:40 (v/v) Matrigel in PBS. Every 2 days the monolayer medium was replaced and the transepithelial electrical resistance (TEER) was measured with a STX4 electrode (EVOM^3^; World Precision Instruments, Berlin, Germany). The differentiation was started at cultivation day 8 by changing the medium to differentiation medium (S3 Table). Differentiation medium was replaced daily and the TEER measured. Experiments were conducted on the 10^th^ day of cultivation.

### RNA isolation and reverse transcription

On the 10^th^ day of cell cultivation, the cells were harvested for analysis. After aspirating the medium, cells grown on transwells were washed with ice-cold PBS and the cellular layer was detached from the membrane with a spatula. For the examination of tight junction proteins, cells were also collected on the 4^th^, 7^th^, and 9^th^ days of cultivation. Samples were centrifuged at 1,000 x *g* for 10 minutes at 4 °C, followed by swift freezing in liquid nitrogen and subsequent storage at -80 °C. The RNA extraction was performed with the RNeasy Plus Mini Kit (Qiagen, Hilden, Germany) according to the manufacturer’s instructions. The thawed cell pellets were resuspended in RLT buffer with 1:100 (v/v) β-mercaptoethanol (Sigma-Aldrich) and the cells were homogenized by ultrasound. Following the removal of gDNA and subsequent precipitation with 70% (v/v) ethanol, the RNA was bound to a column, washed with 700 µl RW1 buffer, centrifuged for 1 min at 16,000 x *g*, than washed two times with 500 µl RPE buffer and centrifuged for 1 min at 16,000 x *g*. The RNA was eluted from the column with 30 µl RNase free water by centrifugation for 1 min at 9,500 x *g*. The concentration of the isolated RNA was determined by spectrophotometric measurements using the NanoDrop™ One (Thermo Scientific™). The cDNA synthesis was performed via reverse transcription using TaqMan™ Reverse Transcription Reagents Kit (Applied Biosystems, Roche Molecular System, Darmstadt, Germany).

### Quantitative real-time PCR

To determine mRNA expression of the target genes, quantitative realtime PCR was used. The gene expression of the genes in S4 Table was determined using the SYBR Green® PCR assay as previously described by Elfers et al. (13). For the targeted gene MUC4 the annealing temperature was 63 °C and for ENaC alpha and NHE3 the annealing temperature was 65 °C for all other genes the annealing temperature was 60 °C. The gene expression of the target genes (S5 Table) were determined by TaqMan®. The reaction mixture contained TaqMan™ Gene Expression Master Mix (Applied Biosystems), 16 ng reverse transcribed cDNA, 300 nM of the specific primers and 100 nM of specific probe at a total volume of 20 µl. For each gene, a negative control and a standard series, which is a dilution series of the amplification product, were used to determine the primer efficiency. The amplification and detection was performed with the real-time PCR cycler Bio-Rad CFX96™ (BIO-RAD Laboratories, INC., Hercules, USA). The housekeeping genes of the 60S ribosomal protein L4 (RPL4) and the 40S ribosomal protein S18 (RPS18) were used to normalise the gene expression of the genes of interest. The relative gene expression was calculated with the Pfaffl (14) method and the mean of the two houskeeping genes.

### Ussing chamber experiments – transport physiology

In order to investigate epthelial transport characteristics, 2D monolayers were mounted in Ussing chambers at the end of differentiation as described by Hoffmann et al. (12). After an initial period of 5 min, the tissues were set to short-circuit-conditions for another 15 min to allow equilibration. The I_sc_ and Rt were monitored every 6 seconds for the whole experiment. The 2D culture was than challenged with one of the substances (Table 1). Furthermore, preincubation with aldosterone for 3 h, 8 h and 48 h before Ussing chamber experiments was performed. The epithelia and the control, which had not been preincubated with aldosterone, were subsequently incubated in the Ussing chamber with 100 µM amiloride. Except for forskolin and aldosterone dissolved in DMSO, all other substances are diluted in aqua destillata. Serosal addition of mannitol after the addition of glucose was carried out to ensure osmotic stability. For analysis, the basal values of I_sc_ determined in the Ussing chamber experiments were the mean calculated from the last ten values prior to the addition of an additive. The changes in short-circuit currents (ΔI_sc_) are the difference between the maximum or minimum value after the addition of each substance and the the basal value.

**Table 1:** Application for Ussing chamber experiments (all chemicals were obtained from Sigma-Aldrich, Darmstadt, Germany).

| Substance | Concentration | application | Incubation time |
| --- | --- | --- | --- |
| **DMSO** | 1:1000 | serosal | 15 min |
| **glucose** | 10 mM | mucosal | 15 min |
| **mannitol** | 10 mM | serosal |  |
| **carbachol** | 10 μM | serosal | 15 min |
| **forskolin** | 10 μM | serosal | 15 min |
| **amiloride** | 50 μM | mucosal | 15 min |
| | 100 μM | | |
| | 500 μM | | |
|  | 1 mM |  |  |
| **aldosterone** | 3 nM, | serosal | Preincubation for<br>3 h, 8 h or 48 h |
|  | 1 nM, |  |  |
|  | 0.3 nM |  |  |

### Scanning electron Microscopy

Electron microscopy sample preparation was performed in a similar way as previously described (15). In brief, cells on transwells were fixed in a 0.1 M EM Hepes buffer with 5% formaldehyde and 2% glutaraldehyde and washed twice to remove aldehyde residues. Dehydration was carried out in a gradient series of EtOH on ice (10 %, 30 %, 50 %, 70 %, 90 %), each step for 10 min and two steps (100 % EtOH) at room temperature. Ethanol was used to prevent damage of the transwell membrane. Afterwards, critical point drying (CPD300 from Leica) and sputter coating with gold palladium (SCD 500 from Bal-Tec) was performed. Samples were visualized with a field-emission scanning electron microscope Merlin (Zeiss) at an acceleration voltage of 5 kV and making use of both, an Everhart Thornley HESE2 and an inlens SE detector.

### Data analysis and statistics

TEER values of the 8^th^ day of cultivation were set to 100% and the others calculated accordingly. The statistical analyses were performed using Prism version 9.0.0 (GraphPad, San Diego, USA). The normal distribution of all data was tested with the D’Agostino & Pearson test. Comparison between mean values on the 10^th^ cultivation day and native colon tissue was conducted using the unpaired t-test. The one-way ANOVA was employed to compare various cultivation times followed by Tukey’s test for post-hoc analysis to correct for multiple comparisons. The paired t-test was employed to compare the mean basal and maximum/minimum I_sc_ values in the presence of a normal distribution. When the data did not follow a normal distribution, the Wilcoxon test was used for the same purpose. Differences were considered statistically significant when p value <0.05. Within figure legends, the number of biological experiments is indicated by the symbol “n.”

## Results

### TEER progression shows robust cellular monolayer integrity

Measurement of TEER provides an indication of the integrity of the cellular layer. Monolayers are confluent on day 8 of cultivation and the differentiation was started by changing the cultivation medium from monolayer medium to differentiation medium. Therefore, the electric resistance was set to 100 % on this day. The progression of the TEER values showed an increase until the 8^th^ day of cultivation and formed a plateau during the differentiation period (Figure 1).

**Figure 1:**
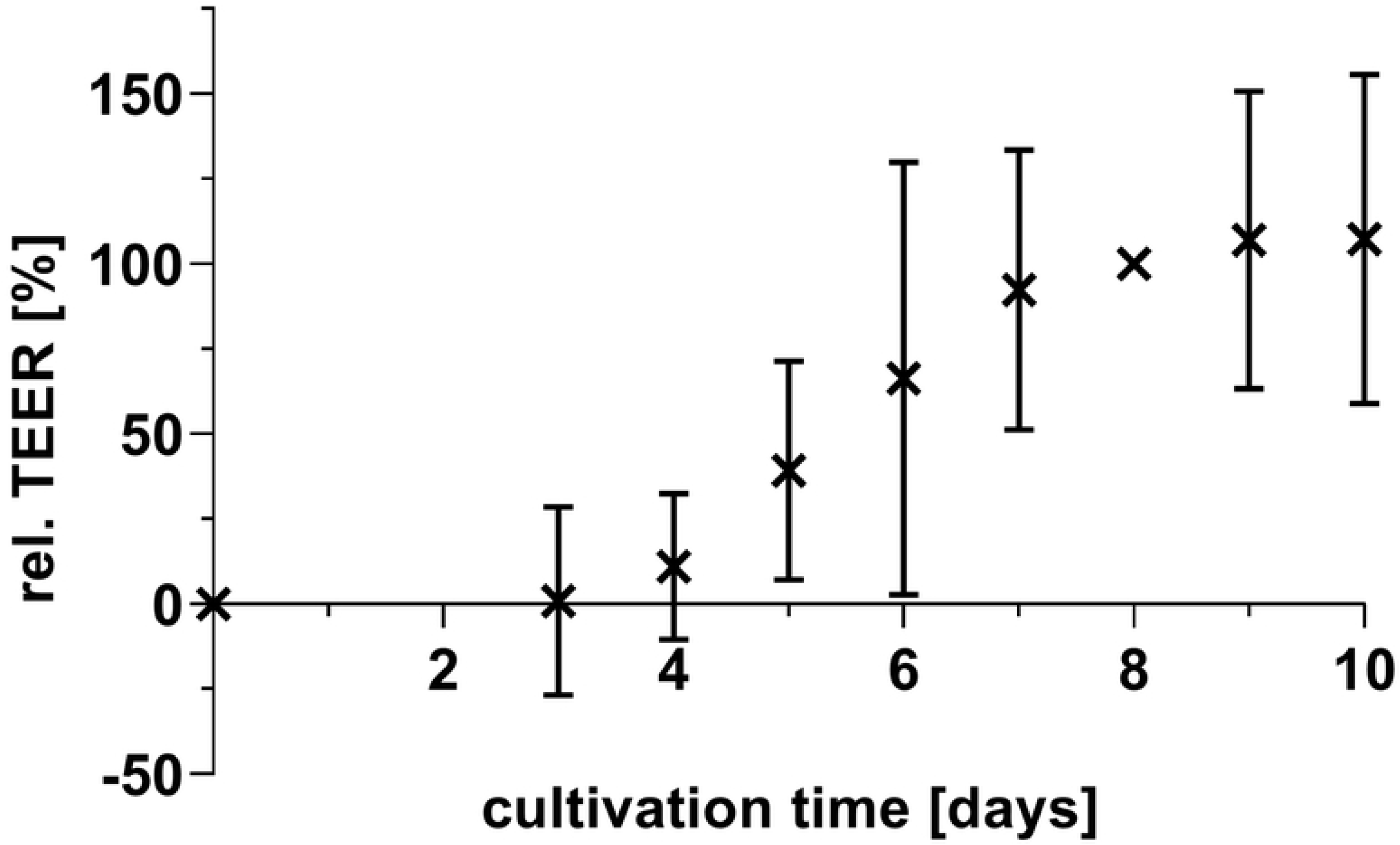
Change of the transepithelial electrical resistance (TEER) during the 2D cultivation of the porcine colonic organoids. The organoids were cultivated on transwell inserts (0.4 µm pore diameter). Displayed is the mean ± SD, n = 11.

### Scanning electron microscopy reveals different cell types

Electron microscopy revealed an intact monolayer of 2D organoids at day 10 of culture. The monolayer was characterized by the development of well-defined microvilli at the apical surface. (Figure 2 A & B). The goblet cells are indentified by multiple orifices, which arise from the fusion of mucin-containing granules (**Error! Reference source not found.** A & B). This process has resulted in the formation of a mucus layer that covers the epithelium (**Error! Reference source not found.** B).

**Figure 2:**
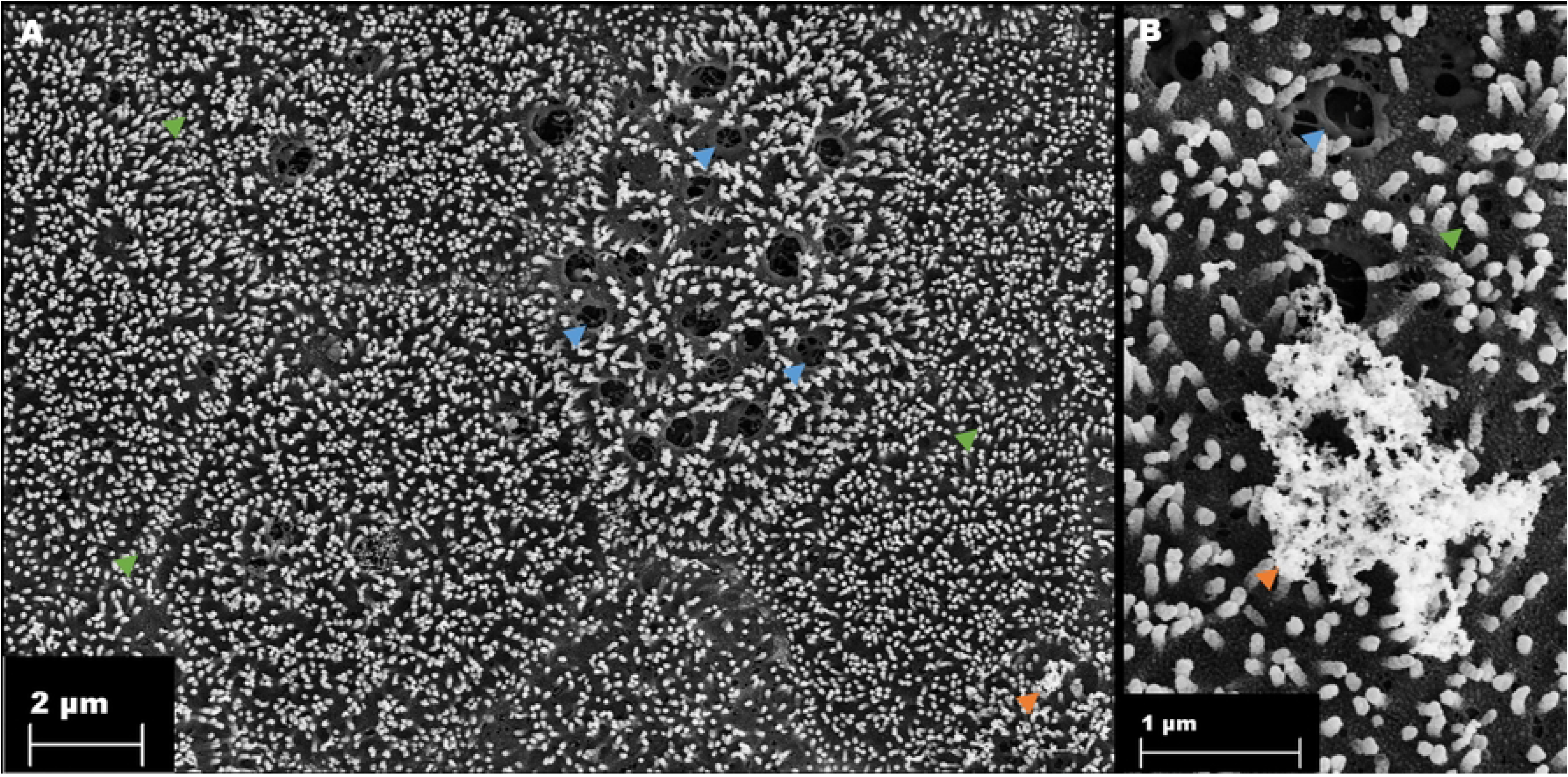
Visualization of microvilli, goblet cells and mucus in the 2D culture of porcine colonic organoids. Green arrows: microvilli; blue arrows: mucin-containing granules; orange arrows: mucus

### RT-PCR demonstrates gene expression concordance between 2D organoids and native tissue

The gene expression of target genes was determined in 2D cultures of organoids and compared with gene expression in native porcine colon tissue. The focus was on key genes that are central to intestinal function, particularly those that control absorption and secretion and are critical for optimal colonic function. Genes involved in the transport of Na^+^, including ENaC and NHE, exhibited no differential expression in the organoid model when compared to native tissue. This pattern was similarly observed for genes for Cl^-^ secretion (CFTR and CaCC), and the gene encoding for the uptake of glucose (SGLT1). Moreover, the expression of the mucin genes (MUC) demonstrated no significant difference between the 2D organoid culture and native tissue (Figure 3).

**Figure 3:**
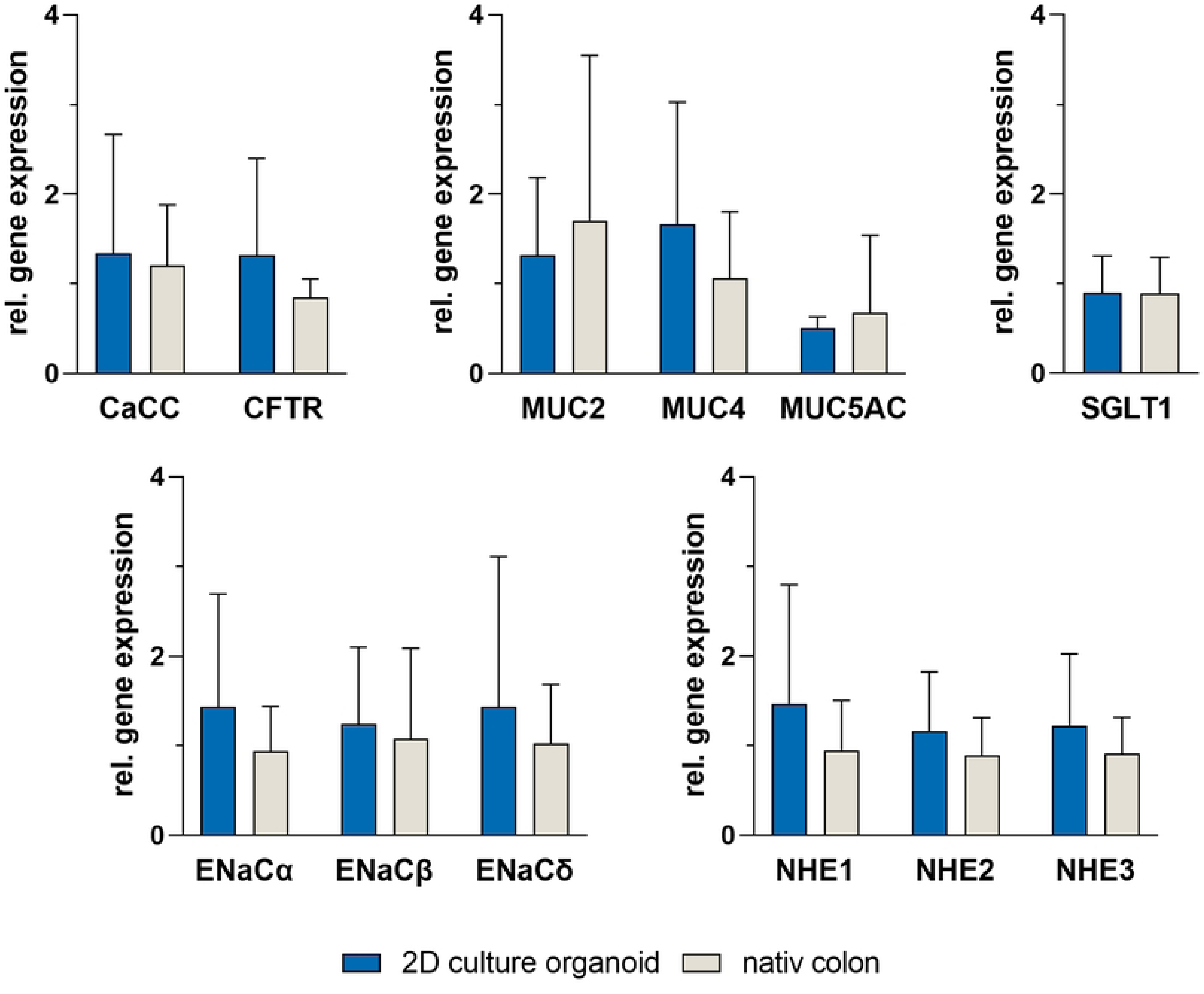
Gene expression of calcium dependent chloride channel (CaCC), cystic fibrosis transmembrane conductance regulator (CFTR), the domains of epithelial sodium channel (ENaC α, ENaC β, ENaC γ, ENaC δ), mucin 2, 4 and 5AC (MUC2, MUC4, MUC5AC), the sodium-potassium-exchanger (NHE1, NHE2, NHE3) and sodium/glucose cotransporter 1 (SGLT1) in 2D cultures of porcine colonic organoids in comparison to native porcine tissue. Ribosomal protein L4 (RPL4) and ribosomal protein S18 (RPS18) were used as reference genes and for normalization. Values shown: mean ± SD, n=4, unpaired t-test was performed with no significant differences.

### Gene expression of tight junction proteins during cultivation is constitutive

The development of tight junction protein gene expression was examined on 4^th^, 7^th^, 9^th^ and 10^th^ day of cultivation. CLDN3 showed no difference in gene expression during the cultivation period on 4^th^, 7^th^, 9^th^ and 10^th^ day. The same was true for CLDN2 with a higher variance on 4^th^ day. The other transmembrane tight junction protein OCLN also showed no significant differences over the cultivation period, but a slightly higher expression on the last day of cultivation (10^th^ day). The tight junction associated protein ZO-1 showed no differences in gene expression over the cultivation period and compared to native colonic tissue (Figure 4).

**Figure 4:**
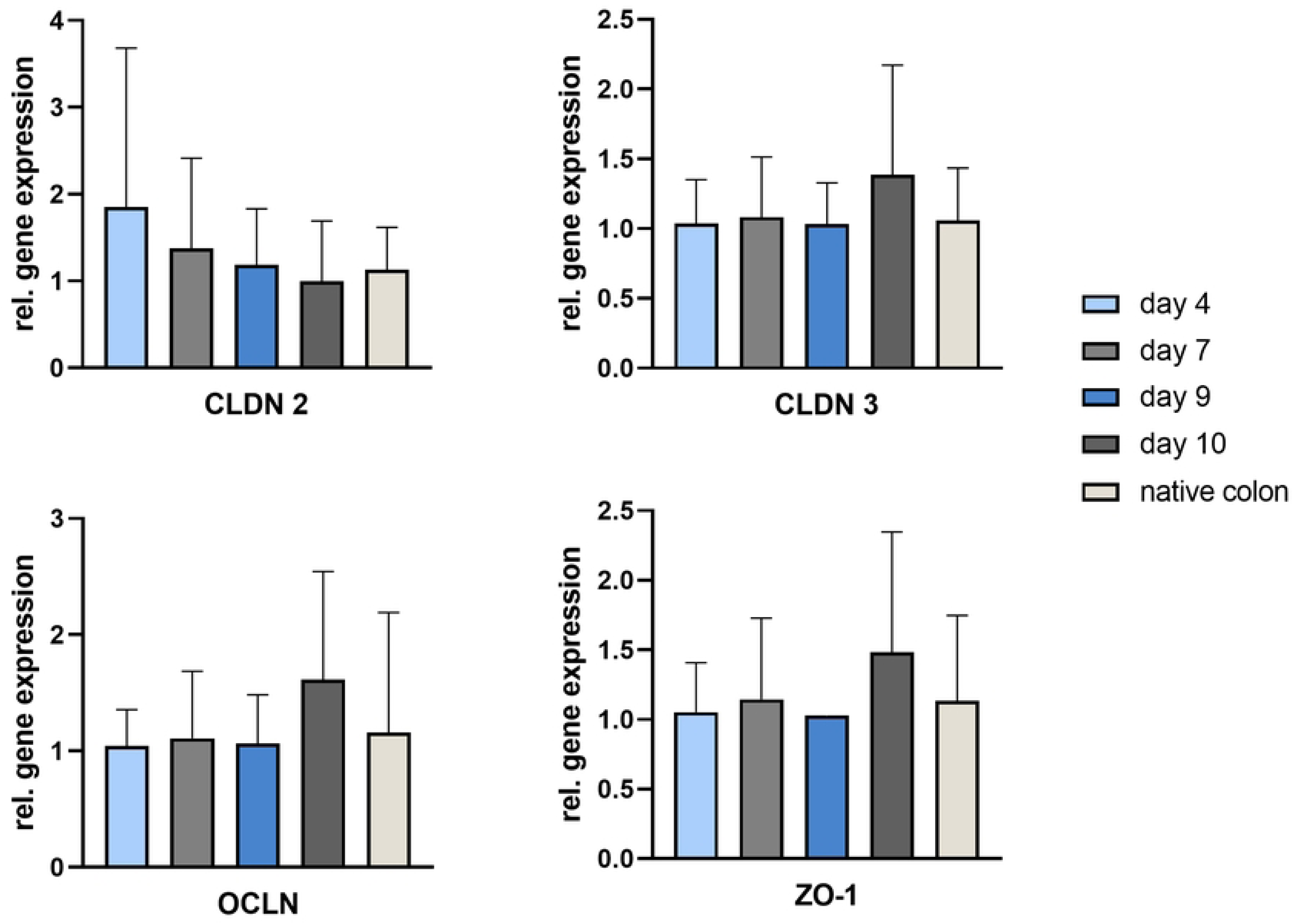
Relative gene expression of the tight junction proteins claudin 2 (CLDN 2), claudin 3 (CLDN 3), occludin (OCLN) and zonula occludens 1 (ZO-1) in 2D-culture of porcine colonic organoids on different cultivation days (day 4, day 7, day 9 and day 10) and native porcine colon. The reference genes ribosomal protein L4 (RPL4) and ribosomal protein S18 (RPS18) were used to normalize the expression. Values shown: mean ± SD, n=4, One-way ANOVA was performed as a statistical test with no significant differences.

### Ussing chamber studies show physiological transport capacity of 2D-organoids similar to native tissues

The addition of DMSO, glucose, forskolin and carbachol was used to alter the I_sc_ in Ussing chamber experiments (Figure 5). The addition of DMSO (0.1 %), which was used as a solvent control, showed no effect to the I_sc_. The same was observed after the addition of carbachol (10^-5^ mol‧l^-1^), which stimulates the Ca^2+^-dependent Cl^-^ secretion. A significant increase in I_sc_ was followed after the addition of glucose (10^-2^ mol‧l^-1^), which was added to test the functionality of the SGLT1, which cotransports Na^+^ with glucose. Forskolin (10^-5^ mol‧l^-1^) activates the cAMP-dependent Cl^-^ secretion through CFTR resulting in a significant increase in I_sc_ after serosal addition (Figure 5).

**Figure 5:**
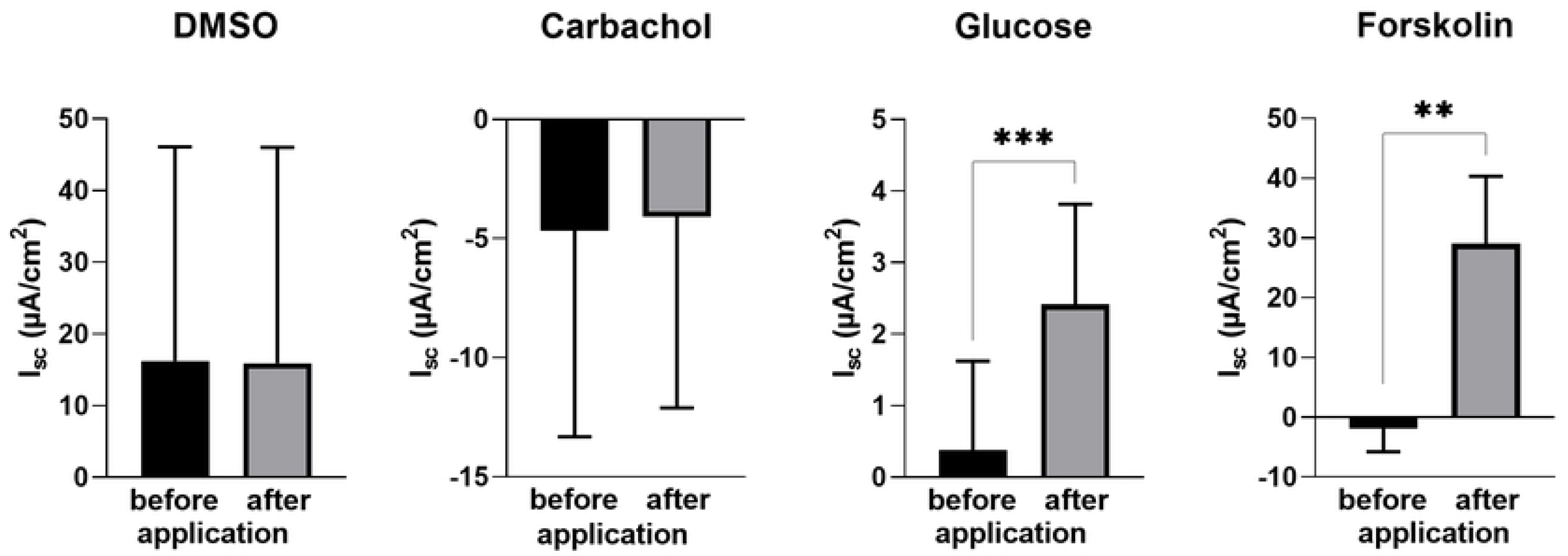
Basal and maximal I_sc_ values of 2D culture of organoids after addition of DMSO, glucose, forskolin and carbachol using Ussing chamber experiments. Values shown: mean ± SD, n=4. For DMSO a Wilcoxon test and for the other a paired t-test was performed: **p<0.01, ***p<0.001.

Amiloride influences the epithelial sodium channel in a dose-dependent manner leading to a significant decrease at 100 µM and 500 µM (Figure 6A). This effect could not be observed with 50 µM or 1 mM amiloride. The addition of forskolin after amiloride led to a significant increase in I_sc_, but showed no concentration-dependent effects (Figure 6B).

**Figure 6:**
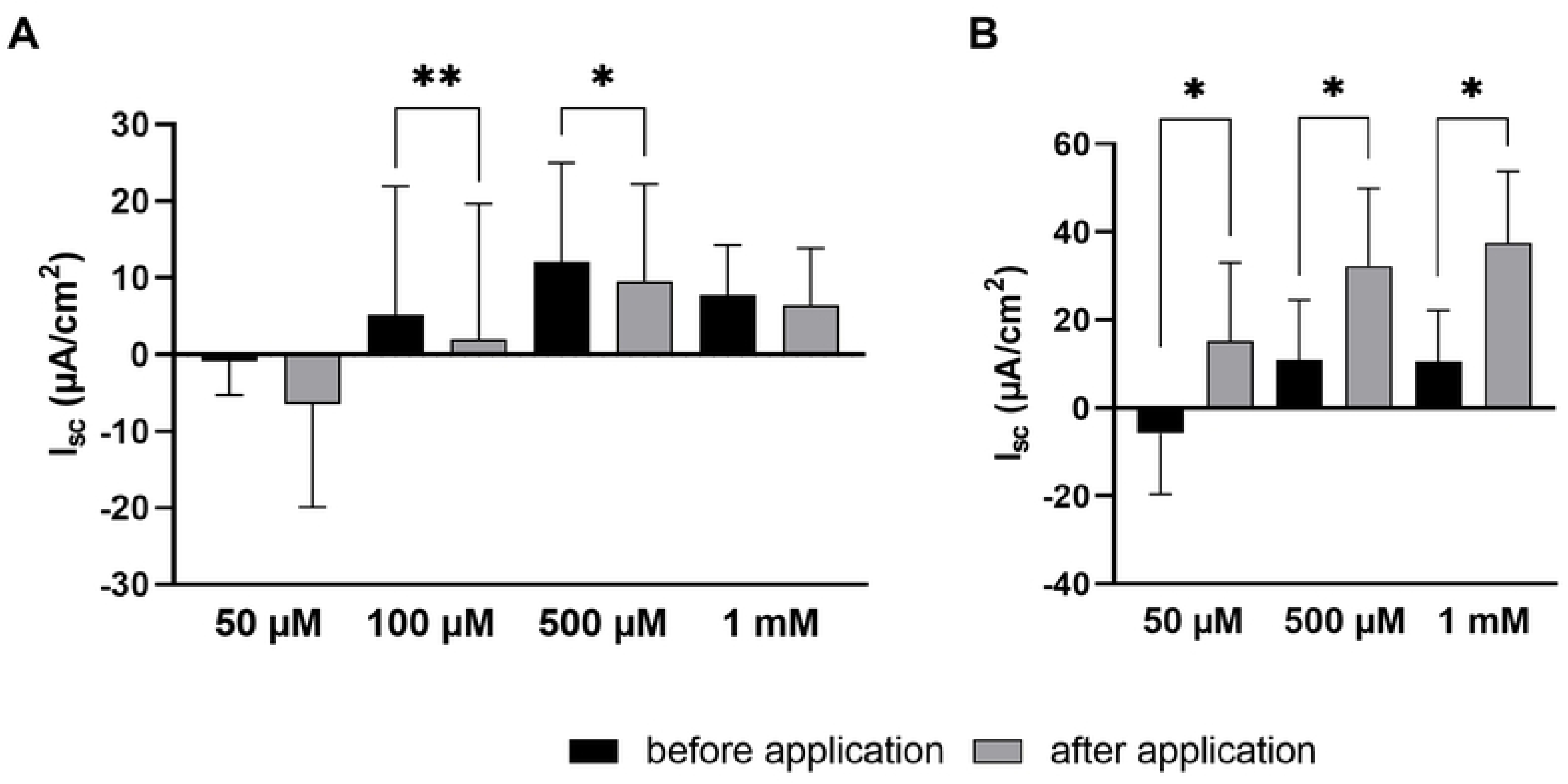
Comparison of the I_sc_ in 2D culture of colonic organoids with different concentrations of amiloride. A: Basal and minimum I_sc_ values after addition of 50 µM, 100 µM, 500 µM and 1 mM amiloride. B: Effect of 10 µM forskolin after the addition of amiloride. The forskolin was added 15 min after amiloride to the chamber. Values shown: mean ± SD, n=4, paired t-test. * p<0.05, ** p<0.01.

After the preincubation period with aldosterone, which activates apical Na^+^ channels, the monolayer was challenged with amiloride in the Ussing chamber. The basal I_sc_ did not undergo a notable alteration when 0.3 nM aldosterone was present. When the two higher concentration were introduced, the dispersion of the basal values increased, yet this did not result in a significant rise in basal I_sc_ (Figure 7). Notably, incubation with 1 nM aldosterone resulted in the smallest reduction in ΔI_sc_. Preincubation showed significant concentration effects (Figure 8). Conversely, incubation with 0.3 nM aldosterone did not yield a significant change in ΔI_sc_ for the 3, 8, or 48-hour preincubation periods (Figure 8). Significant alterations, however, were observed for the concentration and incubation time.

**Figure 7:**
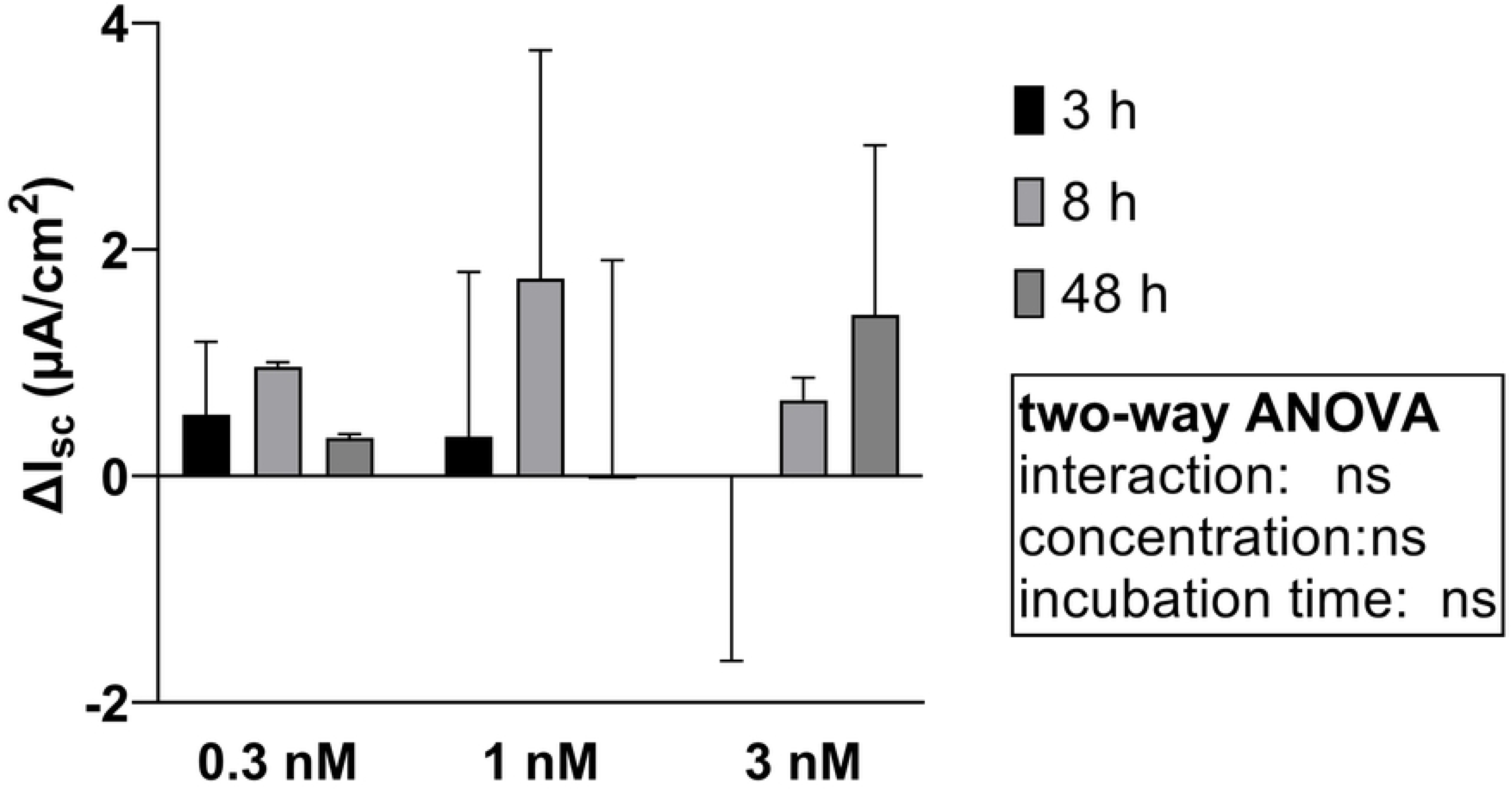
Comparison of the basal I_sc_ in 2D culture of colonic organoids, which were incubated with 0.3 nM, 1 nM or 3 nM aldosterone for 3 h, 8 h or 48 h, while the control was not incubated with any aldosterone. The ΔI_sc_ is the difference between of the aldosterone incubated basal I_sc_ and the control basal I_sc_. Values shown: mean ± SD of four independent experiments, two-way ANOVA with no significant changes

**Figure 8:**
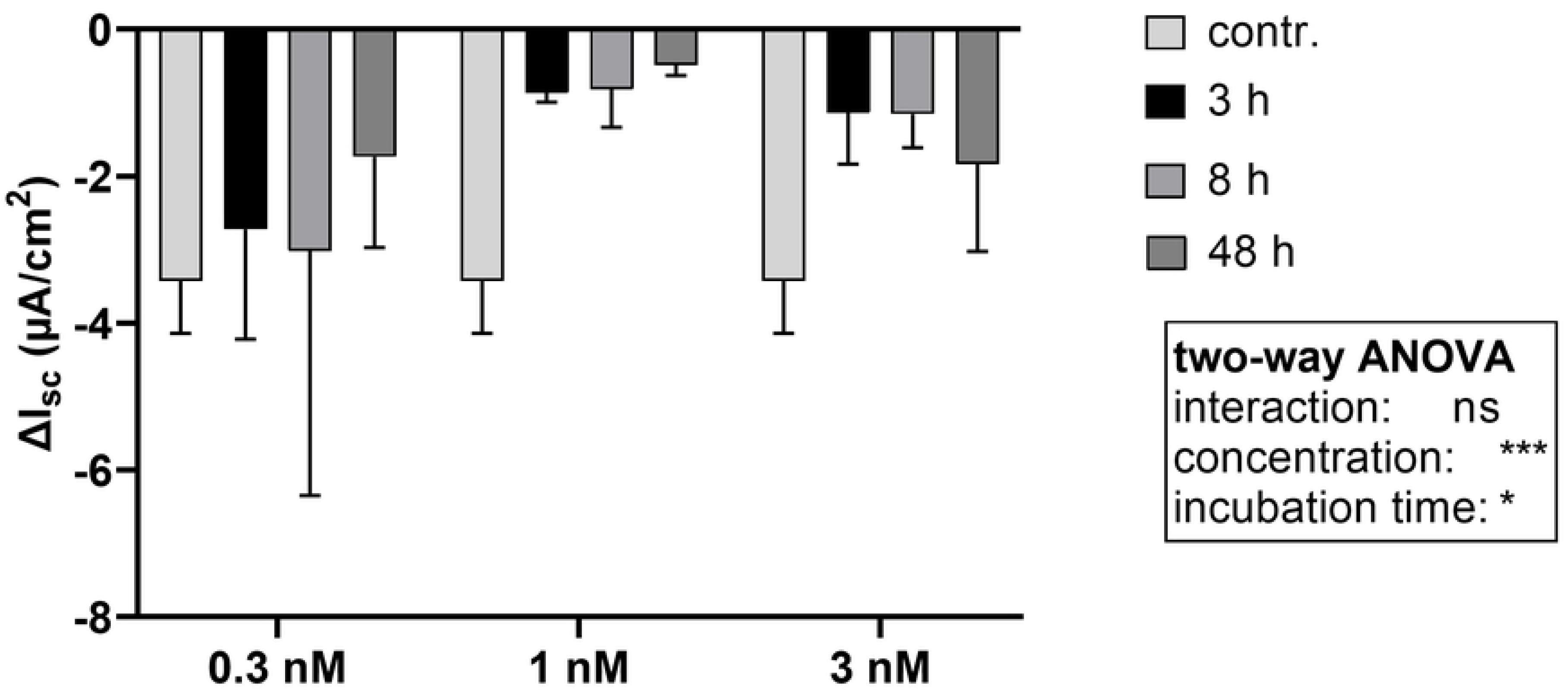
Comparison of the ΔI_sc_ in 2D culture of colonic organoids, which were incubated with 0.3 nM, 1 nM or 3 nM aldosterone for 3 h, 8 h or 48 h, while the control (contr.) was not incubated with any aldosterone. The ΔI_sc_ were calculated from the basal I_sc_ and the minimal I_sc_ after the addition of 100 µM amiloride. Values shown: mean ± SD, n=4, two-way ANOVA. ns > 0.05, * p<0.05, *** p<0.001

## Discussion

### Barrier function

The colonic epithelium maintains selective barrier function, regulating the passage of nutrients, ions and water while safeguarding against luminal pathogens and mechanical stress. Electron microscopy imaging revealed the presence of microvilli on the polarised monolayer and identified goblet cells secreting mucus by exocytosis. These features are consistent with those of the colonic epithelium (16, 17). One part of the barrier function is the mucus, which is composed of mucins. The mucins are divided into two groups, with MUC2 serving as the major structural component. In Addition, MUC4, a membrane-associated mucin, is present in the colon (18). The expression of both MUC2 and MUC4 has been observed in native colon epithelium and in two-dimensional organoid cultures. In contrast, MUC5AC is not typically found in the colon but is more prevalent in the respiratory tract (19), which explains its lower gene expression levels compared to MUC2 and MUC4. However, an increase in MUC5AC secretion may occur in the context of gastrointestinal infections. To gain further insight and utilise the model for the study of pathophysiology, an analysis of gene expression was conducted. The organoids grown in 2D showed increasing TEER until day 8, which then plateaued until the end of the cultivation period. TEER measurement is a method used to monitor the integrity and barrier function of the monolayer (20). The formation of the monolayer resulted in an increase in TEER, and its stability was observed during the last two days of cultivation.

### Functionality of transporters and channels

The barrier function is mediated by tight junction proteins. The tight junction is formed by transmembrane proteins, such as claudins and occludin, together with tight junction-associated proteins that link the transmembrane protein to the actin cytoskeleton. The expression of claudins is organ-specific, and in the colon epithelium *CLDN2*, *CLDN3*, *OCLN*, and *ZO-1* are expressed (2, 21, 22). No differences in the gene expression of these proteins were observed during the cultivation period. The formation of tight junctions between the cells results in an increase in TEER values, which can be observed during the formation of the monolayer in two-dimensional (2D) culture. Since the gene expression of the observed tight junction proteins remains unchanged, we assume that the tight junction proteins are formed constitutively during cultivation and are assembled as a tight junction when the cells grow together to form a monolayer. Consequently, the formation of cell contacts in the monolayer leads to an increase in the TEER value.

The barrier function is of considerable importance in the prevention of pathogen and other microbial invasion. Furthermore, the epithelium plays a role in the absorption of nutrients and the transport of water and ions, which also impacts the rheological properties of the mucus. The function of transporters and channels that play a pivotal role in the colon was examined. The expression of *CaCC* and *CFTR*, both of which secrete Cl^-^ ions to maintain cellular homeostasis, exhibited no statistically significant differences between the model and native tissue. The same was true for *ENaC* as well as the sodium-glucose transporter *SGLT1*. Despite *SGLT1* is known to be expressed to a major extent in the small intestine, some studies have demonstrated that the *SGLT1* is also expressed in the large intestine in various animal species (23-25). It can be concluded that *SGLT1* expression is not a limitation of this model; rather, it is a reflection of its physiological relevance in the large intestine. The observed expression of these transporters and channels in the 2D organoids and native pig tissue indicates that the 2D culture of colonic organoids may serve as a model to mimic the native colonic epithelium.

To examine the functionality of these channels and transporters we applied Ussing chamber experiments. The addition of glucose to the mucosal layer resulted in an increase in I_sc_, which was can be explained by cotransport of glucose and sodium through SGLT1 (26). This response to glucose differed from that observed in native pocine colonic epithelium, where no increase in I_sc_ was measured despite *SGLT1* expression. It is possible that the discrepancy between the organoids and native tissue was due to the former cultivation with glucose as the only carbon source. Yoshikawa et al. (25) observed an increase in *SGLT1* expression in the colon of germ-free mice, where SCFA were absent as a carbon source. In contrast to the native colon, which primarily utilises SCFA derived from fermentation of fietary fibre for energy production (27), this experimental system relied exclusively on glucose. This resulted in the cells exhibiting atypical behaviour compared to their typical SCFA uptake in the native colon.

Chloride can be secreted by the CaCC and the CFTR and the functionality of these channels can be tested in the Ussing chamber with carbachol or forskolin, respectively. The Cl^-^ secretion in the 2D culture of colonic organoids is driven by the CFTR. Forskolin stimulates the CFTR through an increased cyclic adenosin monophosphate (cAMP) level. Addition of forksolin resulted in an increase of the I_sc_ in the Ussing chamber, which is also the case in native porcine colonic tissue (28, 29). The 2D culture of porcine colonic organoids showed no response to carbachol but an increased I_sc_ after forskolin treatment. The Cl^-^ secretion by the CaCC can be stimulated, in native porcine colonic tissue, with the addition of carbachol through an increase of the intracellular Ca^2+^ concentration (30, 31). In native porcine colon the addition of carbachol led to an increased I_sc_ (28, 32). The different response of native colon tissue and 2D colonic organoids to carbachol showed a limitation of the model, because unlike in the native tissue, no functional CaCC can be assumed in the model. The observed lack of response to carbachol in the 2D organoid model is plausibly due to a discrepancy in protein expression. Although no significant difference in gene expression between 2D organoids and native colonic tissue has been detected, the discrepancy between gene expression and protein functionality may be attributable to post-translational modifications, improper protein localisation, or the detection method’s limited sensitivity. To elucidate this further, it would be beneficial to analyse protein expression in future studies using techniques such as Western blotting or immunohistology, which could provide insights into the presence and functionality of the CaCC protein. Furthermore, the lack of response may be attributable to the Ussing chamber method, which may lack the sensitivity required to detect the carbachol-induced effect. It is established that the response to carbachol in the porcine gastrointestinal tract is less pronounced in comparison to forskolin (28), a trend that is also seen in porcine jejunum organoids (12). In light of these considerations, the absence of a discernible carbachol effect may be attributed to the Ussing method’s inadequate sensitivity. In comparison to native tissue, the weaker response in organoids may be further obscured by the method’s limited resolution, potentially masking subtle changes in chloride secretion. Amiloride has been demonstrated to inhibit the ENaC, which results in a reduction in I_sc_ when measured in Ussing chambers (33, 34). We observed this effect in our 2D culture of colonic organoids. Furthermore, the results indicate a concentration-dependent response. Inagaki et al. (34) also demonstrated a dose-dependent inhibition of ENaC in rat colon. No decrease in I_sc_ was observed for the highest concentration of 1 mM amiloride, which was determined to be too high for the inhibition of ENaC. It is established that amiloride inhibits another Na^+^ entry pathway via the NHE, although higher concentrations of amiloride are required for this effect (35). The NHE is an electroneutral exchanger, a characteristic that prevents its detection using the Ussing chamber technique. In particular, it has been demonstrated that 1 mM amiloride inhibits the activity of NHE (36, 37). A concentration of 1 mM amiloride did not result in a reduction in I_sc_ in the 2D culture of porcine colonic organoids, as the inhibition was not of the ENaC, but rather of the NHE.

Additionally, aldosterone exerts an influence on the ENaC. The introduction of aldosterone results in an augmentation of the number of these channels, thereby leading to an increase in the basal current (38-40). In the present 2D culture, no increased basal current was observed after incubation for 3 h, 8 h and 48 h. Following the addition of the antagonist amiloride, a smaller decrease in I_sc_ was observed than in the control, which was not incubated with aldosterone and showed a greater decrease in I_sc_.

The aim of this study was to develop an in vitro model to study functional properties of the porcine colon. In order for the colonic epithelium to be classified as such, it is necessary for it to form a dense monolayer. This was confirmed in the 2D culture of porcine colon organoids through TEER measurements and microscopic imaging. Furthermore, electron microscopy revealed that this 2D monolayer is composed of various cell types. Gene expression in our model does not differ from native tissue, allowing accurate comparisons of gene expression. Functionally, the model closely mimics native porcine colon epithelium, providing a robust platform for multiple comparisons with native tissue. This model supports the 3R-principles by potentially replacing animal experiments in studies of colonic epithelial physiology, infection and pharmacological research.

## Acknowledgement

Thanks to the Federal Ministry of Food and Agriculture (BLE# 28N-2-071-00) for funding.

## Supporting information

**S1 Table:** Compostion of organoid medium

**S2 Table:** Composition of Monolayer medium

**S3 Table:** Composition of Differentiation medium

**S4 Table:** SYBR Green primers for SYBR Green® assay

**S5 Table:** TaqMan primers for TaqMan® assay

## References

1. Hornbuckle WE, Simpson KW, Tennant BC. Gastrointestinal function. Clinical Biochemistry of Domestic Animals. 2008.

2. Kuo WT, Odenwald MA, Turner JR, Zuo L. Tight junction proteins occludin and ZO-1 as regulators of epithelial proliferation and survival. Ann N Y Acad Sci. 2022;1514(1):21–33.

3. Tsukita S, Furuse M, Itoh M. Multifunctional strands in tight junctions. Nature Reviews Molecular Cell Biology. 2001;2(4):285–93.

4. Pelaseyed T, Bergström JH, Gustafsson JK, Ermund A, Birchenough GM, Schütte A, van der Post S, Svensson F, Rodríguez-Piñeiro AM, Nyström EE, Wising C, Johansson ME, Hansson GC. The mucus and mucins of the goblet cells and enterocytes provide the first defense line of the gastrointestinal tract and interact with the immune system. Immunol Rev. 2014;260(1):8–20.

5. Ziegler A, Gonzalez L, Blikslager A. Large Animal Models: The Key to Translational Discovery in Digestive Disease Research. Cell Mol Gastroenterol Hepatol. 2016;2(6):716–24.

6. Russell WMS, Burch RL. The principles of humane experimental technique: Methuen; 1959.

7. Sellers RS, Morton D. The Colon:From Banal to Brilliant. Toxicologic Pathology. 2014;42(1):67–81.

8. Roura E, Depoortere I, Navarro M. Review: Chemosensing of nutrients and non-nutrients in the human and porcine gastrointestinal tract. Animal. 2019;13(11):2714–26.

9. Sato T, Vries RG, Snippert HJ, van de Wetering M, Barker N, Stange DE, van Es JH, Abo A, Kujala P, Peters PJ, Clevers H. Single Lgr5 stem cells build crypt-villus structures in vitro without a mesenchymal niche. Nature. 2009;459(7244):262-5.

10. Zhao Z, Chen X, Dowbaj AM, Sljukic A, Bratlie K, Lin L, Fong ELS, Balachander GM, Chen Z, Soragni A, Huch M, Zeng YA, Wang Q, Yu H. Organoids. Nature Reviews Methods Primers. 2022;2(1):94.

11. Al Shihabi A, Davarifar A, Nguyen HTL, Tavanaie N, Nelson SD, Yanagawa J, Federman N, Bernthal N, Hornicek F, Soragni A. Personalized chordoma organoids for drug discovery studies. Science Advances. 2022;8(7):eabl3674.

12. Hoffmann P, Schnepel N, Langeheine M, Kunnemann K, Grassl GA, Brehm R, Seeger B, Mazzuoli-Weber G, Breves G. Intestinal organoid-based 2D monolayers mimic physiological and pathophysiological properties of the pig intestine. PLoS One. 2021;16(8):e0256143.

13. Elfers K, Marr I, Wilkens MR, Breves G, Langeheine M, Brehm R, Muscher-Banse AS. Expression of Tight Junction Proteins and Cadherin 17 in the Small Intestine of Young Goats Offered a Reduced N and/or Ca Diet. PLoS One. 2016;11(4):e0154311.

14. Pfaffl MW. A new mathematical model for relative quantification in real-time RT-PCR. Nucleic Acids Res. 2001;29(9):e45.

15. Bense S, Witte J, Preusse M, Koska M, Pezoldt L, Droge A, Hartmann O, Musken M, Schulze J, Fiebig T, Bahre H, Felgner S, Pich A, Haussler S. Pseudomonas aeruginosa post-translational responses to elevated c-di-GMP levels. Mol Microbiol. 2022;117(5):1213–26.

16. Barnett AM, Mullaney JA, Hendriks C, Le Borgne L, McNabb WC, Roy NC. Porcine colonoids and enteroids keep the memory of their origin during regeneration. Am J Physiol Cell Physiol. 2021;320(5):C794–C805.

17. Ambrosini YM, Park Y, Jergens AE, Shin W, Min S, Atherly T, Borcherding DC, Jang J, Allenspach K, Mochel JP, Kim HJ. Recapitulation of the accessible interface of biopsy-derived canine intestinal organoids to study epithelial-luminal interactions. PLoS One. 2020;15(4):e0231423.

18. Song C, Chai Z, Chen S, Zhang H, Zhang X, Zhou Y. Intestinal mucus components and secretion mechanisms: what we do and do not know. Exp Mol Med. 2023;55(4):681–91.

19. Ma J, Rubin BK, Voynow JA. Mucins, Mucus, and Goblet Cells. Chest. 2018;154(1):169–76.

20. Srinivasan B, Kolli AR, Esch MB, Abaci HE, Shuler ML, Hickman JJ. TEER measurement techniques for in vitro barrier model systems. Journal of laboratory automation. 2015;20(2):107–26.

21. Tachibana K, Bai L, Sugimura S, Fujioka H, Kishimoto W, Mizuguchi H, Nakase H, Kondoh M. Heterogenous Gene Expression of Bicellular and Tricellular Tight Junction-Sealing Components in the Human Intestinal Tract. Biol Pharm Bull. 2024;47(6):1209–17.

22. Markov AG, Veshnyakova A, Fromm M, Amasheh M, Amasheh S. Segmental expression of claudin proteins correlates with tight junction barrier properties in rat intestine. J Comp Physiol B. 2010;180(4):591–8.

23. Gonzalez LM, Williamson I, Piedrahita JA, Blikslager AT, Magness ST. Cell lineage identification and stem cell culture in a porcine model for the study of intestinal epithelial regeneration. PLoS One. 2013;8(6):e66465.

24. Bindslev N, Hirayama BA, Wright EM. Na/D-Glucose Cotransport and SGLT1 Expression in Hen Colon Correlates with Dietary Na+. Comparative Biochemistry and Physiology Part A: Physiology. 1997;118(2):219–27.

25. Yoshikawa T, Inoue R, Matsumoto M, Yajima T, Ushida K, Iwanaga T. Comparative expression of hexose transporters (SGLT1, GLUT1, GLUT2 and GLUT5) throughout the mouse gastrointestinal tract. Histochemistry and Cell Biology. 2011;135(2):183-94.

26. Wright EM, Hirsch JR, Loo DD, Zampighi GA. Regulation of Na+/glucose cotransporters. The Journal of experimental biology. 1997;200(Pt 2):287–93.

27. Bindelle J, Buldgen A, Wavreille J, Agneessens R, Destain JP, Wathelet B, Leterme P. The source of fermentable carbohydrates influences the in vitro protein synthesis by colonic bacteria isolated from pigs. Animal. 2007;1(8):1126–33.

28. Leonhard-Marek S, Hempe J, Schroeder B, Breves G. Electrophysiological characterization of chloride secretion across the jejunum and colon of pigs as affected by age and weaning. Journal of Comparative Physiology B. 2009;179(7):883–96.

29. Bridges RJ, Rummel W, Simon B. Forskolin induced chloride secretion across the isolated mucosa of rat colon descendens. Naunyn-Schmiedeberg’s Arch Pharmacol. 1983;323(4):355–60.

30. Chandan R, Hildebrand KR, Seybold VS, Soldani G, Brown DR. Cholinergic Neurons and Muscarinic Receptors Regulate Anion Secretion in Pig Distal Jejunum. European Journal of Pharmacology. 1991;193(3):265–73.

31. Dharmsathaphorn K, Pandol SJ. Mechanism of Chloride Secretion Induced by Carbachol in a Colonic Epithelial-Cell Line. J Clin Invest. 1986;77(2):348–54.

32. Pfannkuche H, Mauksch A, Gäbel G. Modulation of electrogenic transport processes in the porcine proximal colon by enteric neurotransmitters. Journal of Animal Physiology and Animal Nutrition. 2011;96(3):482–93.

33. Sariban-Sohraby S, Benos DJ. The amiloride-sensitive sodium channel. Am J Physiol. 1986;250(2 Pt 1):C175-90.

34. Inagaki A, Yamaguchi S, Ishikawa T. Amiloride-sensitive epithelial Na+ channel currents in surface cells of rat rectal colon. Am J Physiol Cell Physiol. 2004;286(2):C380–90.

35. Benos DJ. Amiloride: a molecular probe of sodium transport in tissues and cells. Am J Physiol. 1982;242(3):C131–45.

36. Rabbani I, Siegling-Vlitakis C, Noci B, Martens H. Evidence for NHE3-mediated Na transport in sheep and bovine forestomach. Am J Physiol Regul Integr Comp Physiol. 2011;301(2):R313–9.

37. Martens H, Gäbel G, Strozyk B. Mechanism of electrically silent Na and Cl transport across the rumen epithelium of sheep. Exp Physiol. 1991;76(1):103–14.

38. Amasheh S, Milatz S, Krug SM, Bergs M, Amasheh M, Schulzke JD, Fromm M. Na+ absorption defends from paracellular back-leakage by claudin-8 upregulation. Biochem Biophys Res Commun. 2009;378(1):45–50.

39. Fromm M, Schulzke JD, Hegel U. Control of electrogenic Na+ absorption in rat late distal colon by nanomolar aldosterone added in vitro. American Journal of Physiology-Endocrinology and Metabolism. 1993;264(1):E68–E73.

40. Asher C, Wald H, Rossier BC, Garty H. Aldosterone-induced increase in the abundance of Na+ channel subunits. Am J Physiol. 1996;271(2 Pt 1):C605-11.

